# The fusion gene occurrence spectrum predicts the spatial structure of chromosomes

**DOI:** 10.64898/2026.10.07.757465

**Authors:** Xianzhi Ren, Yueqin Cao, Yongliang Dai, Zhiyuan Huang, Junan yu, Shoushuai Feng, Lingfeng Long, Chentao Yang, Qiong Wang, Hailin Yang

**Author notes:** These authors contributed equally to this paper (Xianzhi Ren, Yueqin Cao, Yongliang Dai).

## Abstract

Chromosome was "self-assembled" into a complex three-dimensional topology in the nucleus, which provided the structural basis for important biological processes such as gene expression regulation and cell differentiation. The current understanding of the three-dimensional organization of the genome is based on high-throughput sequencing and imaging technologies to visualize the conformation of chromatin in its native environment. This study proposes a new research perspective and technical means to use fusion genes to determine the interaction information and three-dimensional structure of chromosomes. This three-dimensional information provided by the fusion gene spectrum not only shows the universality of the spatial layout of chromosomes, but also increases the interpretability of the occurrence of fusion genes. The chromatin conformation capture data of multiple tissues and periods is significantly correlated with the strength of fusion genes. For example, the frequency of trans-interaction between chromosomes 11 and 17 was the highest, and the frequency of fusion gene occurrence corresponding to *KMT2A* on chromosome 11 was the highest. Therefore, we infer that this structural universality is a common feature of the human genome, which provides a theoretical basis and data for predicting chromosome fusion events.

**Significance:** Three-dimensional organization of chromosomes is significant for understanding the biological processes, currently, the understanding of the structure of chromosomes and its underlying mechanisms is still largely limited. Here, we determined the sharing pattern of spatial organization through chromosomal translocation data in malignant tumors of multiple tissues and periods. The three-dimensional structure indicated that chromosome 1 was close to the nucleolus, while chromosome 20 was distributed on the periphery. High density of SD sequences near the fusion gene loci and the small difference in GC content between fusion gene pairs were the key factors promoting the formation of fusion genes. This work reveals the causes of fusion genes at the molecular level and maps the universal three-dimensional spatial structure of chromosomes.

## Introduction

The intricate three-dimensional organization of chromosomes within the nucleus is increasingly recognized as a fundamental layer of genomic regulation, influencing processes such as gene expression, DNA replication, and the maintenance of genome stability [1–5]. This non-random architecture facilitates crucial spatial interactions between regulatory elements and their target genes via loop extrusion model and Enhancer-promoter interaction, often located far apart on the linear genome [6]. Furthermore, the accessibility of chromatin is also a key factor regulating gene expression, mediated by epigenetic modifications and chromatin remodeling complexes [7, 8]. Understanding the principles governing this organization is therefore paramount to deciphering the complexities of cellular function and disease. Concurrently, fusion genes, formed by the joining of two previously distinct genes, have emerged as critical players in various biological contexts, most notably in cancer, where they frequently act as potent oncogenic drivers and represent valuable diagnostic and therapeutic targets.

Current methods for elucidating chromosome spatial structure largely rely on experimental techniques. Chromosome conformation capture and its high-throughput derivative Hi-C, along with their variations, provide valuable insights into the average spatial proximity of genomic loci across large cell populations [9]. Microscopy-based methods, such as fluorescence in situ hybridization, allow for the visualization of specific genomic regions in individual cells [10]. These techniques confirmed that chromosomes in mitotic cells are highly self-condensed to form rods-like structure, then become unfolded and distributed in separate domains in interphase cells [11, 12]. In these domains, chromosomes are further subdivided into megabasse chromatin compartmentation, which is subdivided into sub-megabasse topologically associated domains, and chromatin loops in the more sophisticated kilobase range [13–15]. While these approaches have significantly advanced our understanding, they often present limitations. Hi-C data typically reflects a static, population-averaged view, potentially masking the dynamic and probabilistic nature of chromosome interactions. Microscopy techniques, while offering single-cell resolution, are often limited in the number of loci that can be simultaneously examined [16, 17]. Consequently, there remains a need for complementary, predictive approaches that can shed light on the principles governing chromosome spatial organization.

Balanced chromosomal translocations, inversions, deletions, and tandem repeats, collectively known as structural variants (SVs), alter gene dose, location of regulatory elements, and/or copy number of genomics regions, sources of genomic variation [18]. SVs can drive gene fusion events, which are often implicated in cancer [19]. Emerging evidence suggests a reciprocal relationship between SVs and higher-order chromatin conformation [20, 21]. The spatial arrangement of the genome influences the formation and development of SVs, and conversely, SVs can alter spatial organization at a local or even chromosomal level within the nonlinear genome [22, 23]. Fusion gene databases have been developed to catalog gene fusion events cross-species, cancer type and technologies (ChimerDB 4.0, COSMIC, TCGA). Resources such as FusionGDB and OncoKB further annotate the function and mechanism of the specific fusion gene and guide drug use. Integrating and standardizing data from these databases, can facilitate comprehensive analyses, linking mechanisms to clinical guidelines.

In this study, we collected the data of gene fusion data from blood diseases and solid tumors in different cells across different cell types, time points. We mapped the three-dimensional interactions, analyzed the characteristics of chromosomes and genes, and revealed the molecular mechanism of fusion genes from the segmental duplication level. This paper reveals the spatial structure of chromosomes from a new perspective of fusion genes, providing new insights to elucidate the dynamic interaction between 3D genome structure and SV formation, as well as to develop new directions for disease diagnosis and treatment.

## Result

### Three-dimensional spatial structure of chromosomes

Based on the fused gene data, we utilized simulated annealing algorithm to reconstruct the three-dimensional genomic structure with chromosome banding as the basic unit. During the modeling process, the cell nucleus was approximated as a spherical structure to precisely locate the spatial distribution of each chromosome within the nucleus. To quantitatively describe the distribution characteristics of chromosomes in the radial dimension, the cell nucleus was partitioned into four concentric layer regions based on the actual distance range of the banding to the nuclear center and the spatial distribution of the banding: the core layer (0.019-0.535 from the center), the inner middle layer (0.535-0.918), the outer middle layer (0.918-1.364) and the perinuclear layer (1.364-3.427). These four layers contained approximately 192 prominent bands, accounting for a quarter of the total. A large number of literatures have reported that chromosome 19 tends to be located in the central region of the nucleus due to its high gene density and transcriptional activity [24, 25]. Therefore, we added a weighting coefficient to chromosome 19 to make it closer to the nuclear center position in the model. The results showed that chromosome 21 was located on the periphery of the cell nucleus, and 55.6% of it is in the perinuclear layer. 52.9% of chromosome 19 was located in the core layer, adjacent to the nucleolus. The spatial positions of chromosomes 1, 17 and 22 were close to the nucleolus, while chromosomes 5 and 20 were close to the nuclear membrane. The Y chromosome was mainly distributed in the inner and outer middle layer.

### Tran-chromosome interaction

Trans-chromosomal interactions play a crucial role in hematological diseases and solid tumors via promoting chromosomal translocations, the formation of fusion genes and the activation of oncogenic signals. Fig. 2 showed the trans-chromosome interaction events. It is obvious that chromosome 11 was the most prone to trans-chromosome interaction, and the interaction intensity between chromosome 17 and chromosome 11 was the highest, followed by chromosome 3 and chromosome 1. On the whole, the frequency of trans-interaction of was high in megachromosomes, and the interaction between chromosome 1 was the most frequent, with 895 inter-interactions. The number of interactions between chromosomes 17 and 19 of short chromosomes was also higher, with 599 and 591 interactions occurring respectively.

**Figure 1.**
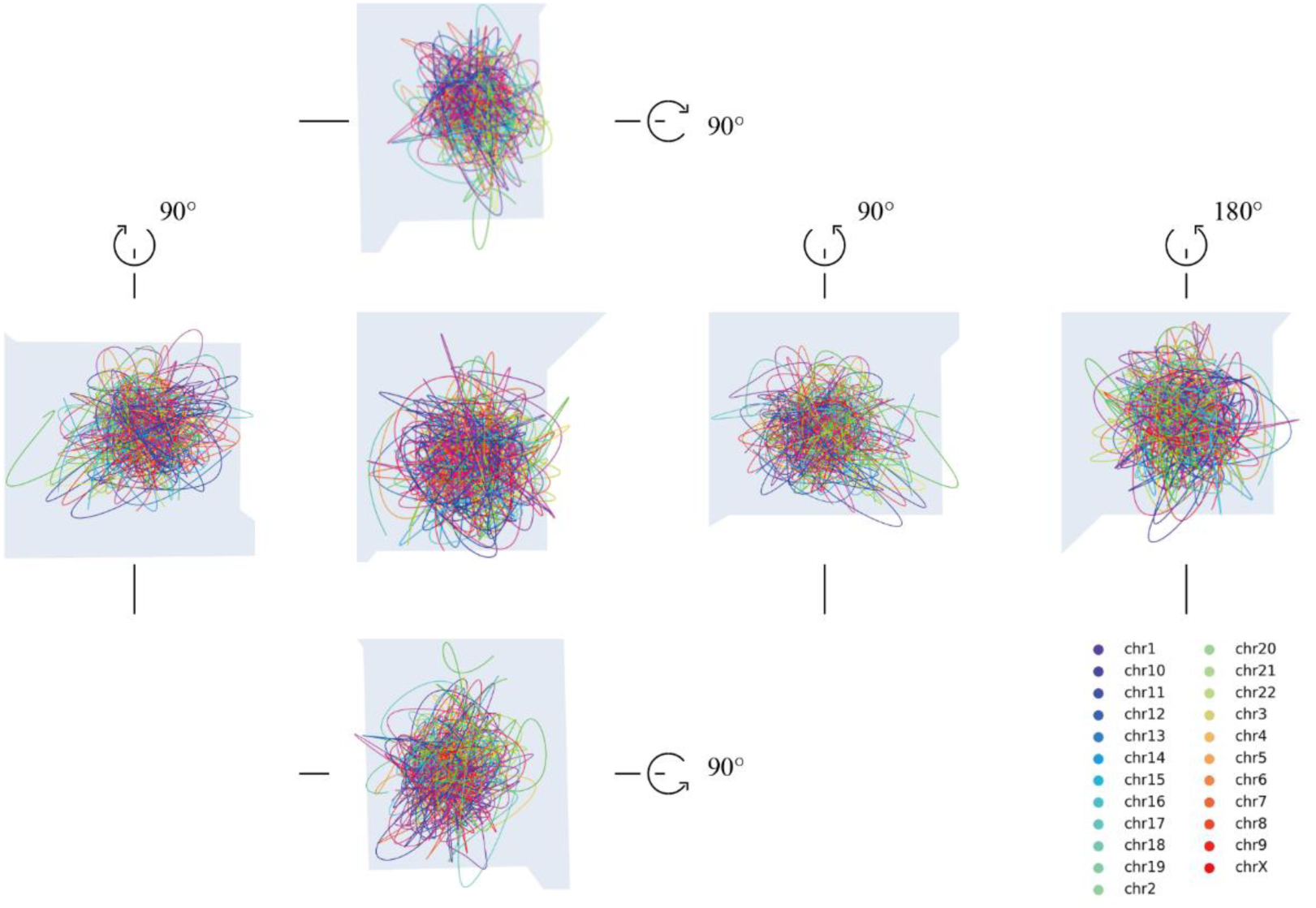
Three-dimensional structures of chromosomes in different dimensions.

**Figure 2.**
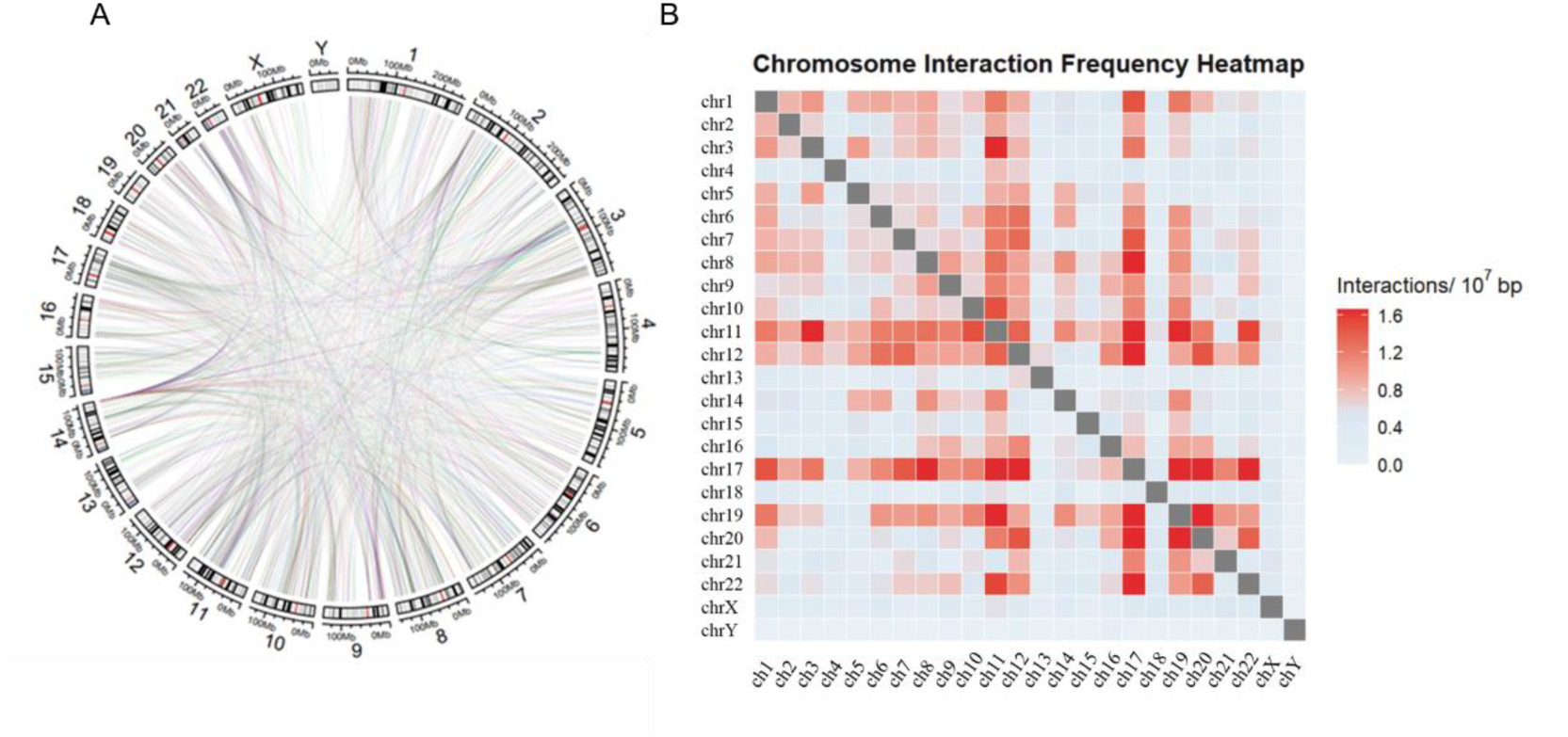
Chromosomal interaction events. A. A circus plot of the trans-interaction of 23 chromosomes. From inner to outside, the circles represent chromosome banding, chromosome size and chromosome number respectively; B. Normalized contact matrix diagram of trans-interaction of 23 chromosomes.

**Figure 3.**
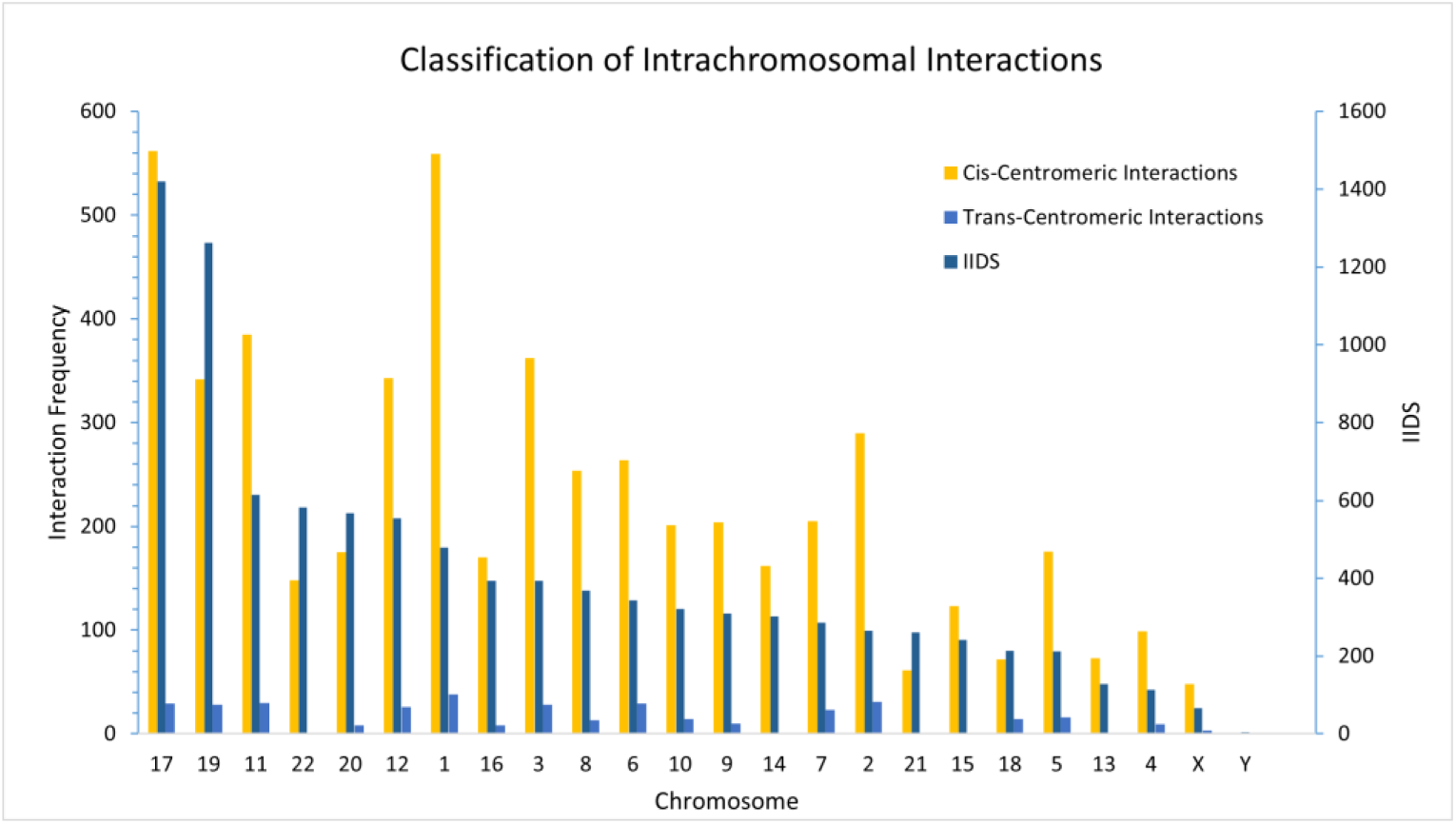
The chromosome interaction events partitions. The figure shows the interaction frequency of different chromosomes (left axis) and the degree of interaction density within chromosomes (right axis).

### Cis-chromosome interaction

To clarify the complexity of chromosomes, we defined the intra-chromosomal interaction density score (IIDS) (Equation 1). The higher the IIDS, the more complex the spatial structure of chromosomes based on internal interactions. The distribution might be more tightly clustered.

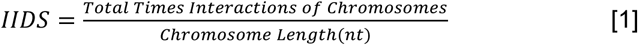

The results indicated that chromosomes 17 and 19 had the highest IIDS, suggesting the complexity of their spatial structure, while chromosomes 4 and 13 had lower IIDS, suggesting that they may be spatially more dispersed in configuration or in a relatively conserved position.

### Chromosome interaction frequency

High-frequency regions of chromosomal interaction are usually spatially adjacent and vulnerable to DNA damage or replication errors, becoming "hotspots" for broken, translocated or fused genes. The interaction frequency scores of chromosomes were shown in Fig. 4, representing chromosomes 1, 2, 3, and 4 (A-D). Overall, the frequency of cis- and trans-interactions in chromosome 1 was the highest, and the frequency of cis-interactions was higher than that of trans-interactions, while the frequency of intra-interaction in chromosome 3 was higher. Most of the genes of chromosome 1 had short length and high interaction frequency, but few interaction times. The internal interaction was very dense, the interaction frequency score and interaction frequency were high, belonging to the fusion intensive genes. The internal interaction of chromosome 2 was mainly characterized by high frequency and low frequency of interaction, indicating that the gene fragments of interaction were long. Chromosome 4, with low interaction frequency and low interaction frequency fraction, was a short gene fraction with loose fusion genes.

**Figure 4.**
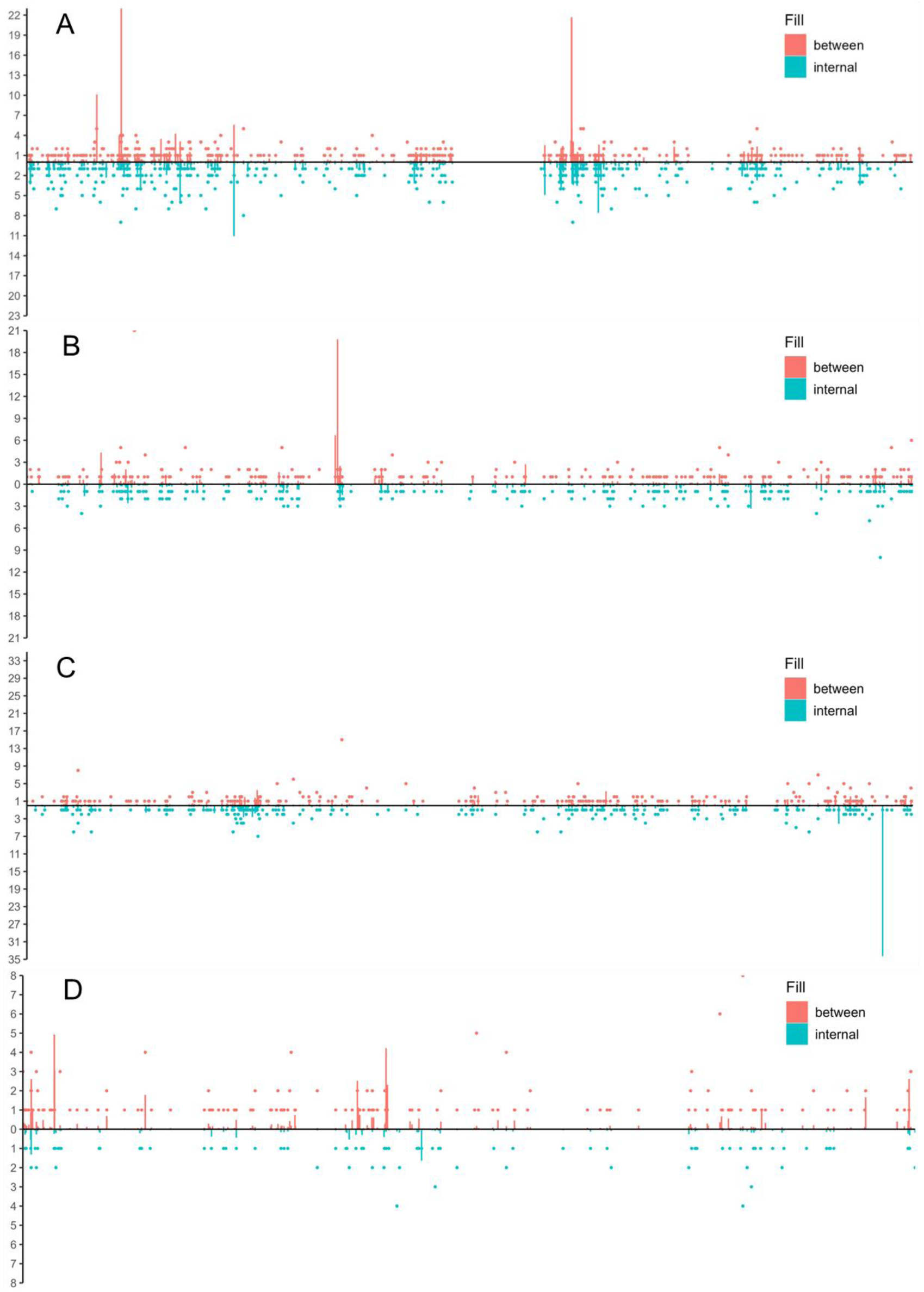
Chromosome interaction frequency fraction. A double Y-axis plot represents the interaction frequency fractions and interaction times of chromosomes 1, 2, 3 and 4 from top to bottom. The horizontal axis of the figure represents the chromosome, different subjects represent different gene positions on the chromosome, the upper part of the vertical axis is the interaction frequency fraction of the interaction between chromosomes (red), the lower part of the vertical axis is the interaction frequency fraction of the interaction within the chromosome (blue), and the dot represents the interaction frequency.

### High interaction frequency gene

Based on the statistical analysis of gene interaction frequency in 23 pairs of chromosomes, this study found a series of genes with high interaction frequency. As shown in Table 1, *KMT2A* on chromosome 11 showed the highest number of inter-and intra-interactions, followed by *ETV6* on chromosome 12 (62), *IGH* on chromosome 14 (53), and *RUNX1* on chromosome 21 (52) with more than 50 inter-interactions. The *MYC* gene on chromosome 8 showed the highest frequency of inter-interaction, with an average of one inter-interaction every 420 bases. This is followed by the *EWSR1* gene on chromosome 22 and the *FUS* gene on chromosome 16. It has been shown that these high-frequency interacting genes were closely related to a variety of hematological malignant diseases, including acute promyelocytic leukemia, acute myeloid leukemia, and myelodysplastic syndrom. These genes have received much attention in hematology research and clinical practice. *RARA* gene abnormality was closely related to the pathogenesis of acute promyelocytic leukemia. The fusion events of *DDX27* gene were mainly concentrated in the field of colorectal cancer, while the abnormality of *FGFR2* gene was associated with the occurrence and development of various malignant tumors.

**Table 1.** Interaction genes with high frequency on chromosomes.

| Gene | Trans-interaction | Cis-<br>interaction | Banding position | Gene Length | Interaction<br>Frequency Score |
| --- | --- | --- | --- | --- | --- |
| *KMT2A* | 101 | 9 | 11q23.3 | 90340 | 12.18 |
| *ETV6* | 62 | 9 | 12p13.2 | 245703 | 2.89 |
| *IGH* | 53 | 4 | 14q32.33 | 1293407 | 0.44 |
| *RUNX1* | 52 | 5 | 21q22.12 | 261501 | 2.18 |
| *EWSR1* | 48 | 3 | 22q12.2 | 32253 | 15.81 |
| *BRAF* | 43 | 36 | 7q34 | 211601 | 3.73 |
| *ALK* | 42 | 27 | 2p23.2 | 728812 | 0.95 |
| *NUP98* | 41 | 5 | 11p15.4 | 122544 | 3.75 |
| *PDGFRB* | 35 | 1 | 5q32 | 42006 | 8.57 |
| *RARA* | 31 | 19 | 17q21.2 | 48463 | 10.32 |
| *PAX5* | 31 | 2 | 9p13.2 | 200999 | 1.64 |
| *RET* | 28 | 10 | 10q11.21 | 53282 | 7.13 |
| *JAK2* | 24 | 8 | 9p24.1 | 145558 | 2.20 |
| *FGFR2* | 23 | 11 | 10q26.13 | 120128 | 2.83 |
| *ROS1* | 23 | 8 | 6q22.1 | 138589 | 2.24 |
| *ABL1* | 23 | 4 | 9q34.12 | 174632 | 1.55 |
| *FGFR1* | 22 | 10 | 8p11.23 | 57492 | 5.57 |
| *FOXP1* | 21 | 4 | 3p13 | 629270 | 0.40 |
| *DDX27* | 18 | 2 | 20q13.13 | 24657 | 8.11 |
| *FUS* | 17 | 2 | 16p11.2 | 14761 | 12.87 |
| *MLLT10* | 16 | 0 | 10p12.31 | 209874 | 0.76 |
| *PVT1* | 16 | 11 | 8q24.21 | 306732 | 0.88 |
| *SIAE* | 16 | 0 | 11q24.2 | 43190 | 3.70 |
| *ARFGEF2* | 15 | 5 | 20q13.13 | 114982 | 1.74 |
| *RAD51B* | 15 | 8 | 14q24.1 | 863317 | 0.27 |
| *BCAS3* | 15 | 28 | 17q23.2 | 714980 | 0.60 |
| *MYC* | 15 | 3 | 8q24.21 | 7517 | 23.95 |

Overall, the interaction frequency score was influenced by both gene length and interaction frequency between genes. Although some genes had more interaction times, their interaction frequency score was relatively low due to the larger gene length. It was worth noting that regardless of the interaction frequency score, these genes played a crucial role in human physiological processes. The interaction frequency score discussed in this study was mainly used to quantify the complexity and density of genes in the spatial structure, providing a new perspective for the identification of core genes. This analysis method based on interaction frequency provided new insights for our understanding of the three-dimensional structure and function of genomes.

### Interaction between *KMT2A*, *NUP98* genes

The interaction between chromosomes was represented by crocs diagram model, taking *NUP98* and *KMT2A* genes on chromosome 11 as examples. Fig. 5 (a) showed that the *KMT2A* gene, located at 11q23.3 and with a length of 90,340 nt, had 101 interactions with different chromosomes, and the interaction frequency score was as high as 12.18. Overall, *KMT2A* had the highest interaction frequency with chromosomes 5 and 17 and 19. The *NUP98* gene, located at 11p15.4 and 122,544 bp in length, interacted with different chromosomes 41 times, or once every 2,988 bases (Fig. 5 (b)). *NUP98* was prone to fuse with the HOX family, accounting for a quarter of the total interaction.

**Figure 5.**
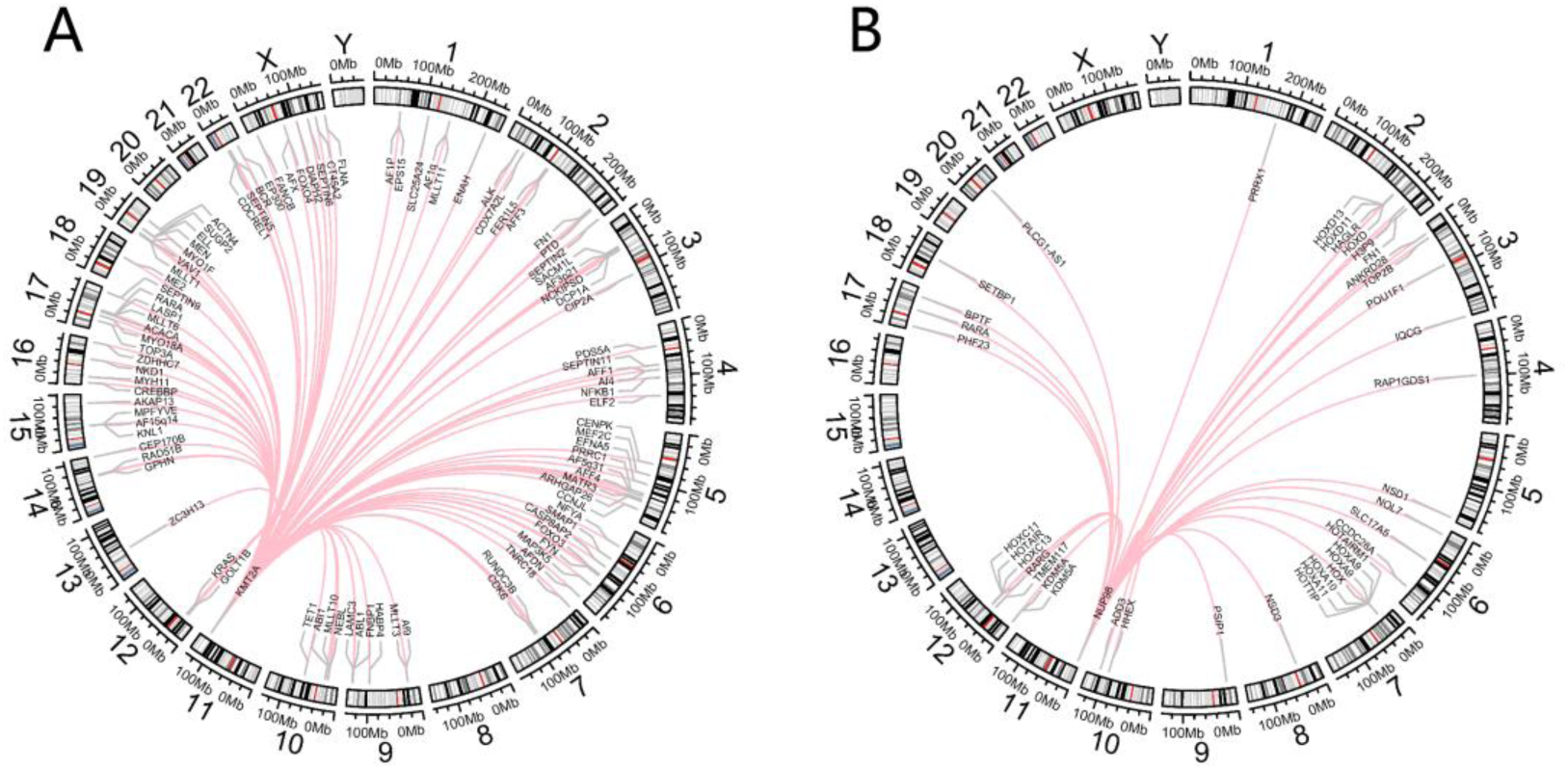
Trans-interaction of *KMT2A* and *NUP98* genes on chromosome 11. A. *KMT2A*; B. *NUP98*.

### Distribution of high frequency interacting genes on chromosomes

As shown in Fig. 6, highly interacting genes (HFIIGs) tend to cluster in regions of chromosomes 11, 19 and 17, indicating that these regions are prone to gene fusion, especially the long arm region of chromosome 17. In contrast, chromosomes 1, 13 and 15 are basically distributed with low interaction genes and are more conserved. This is consistent with the conclusion above that fusion genes tend to occur on hub chromosomes.

**Figure 6.**
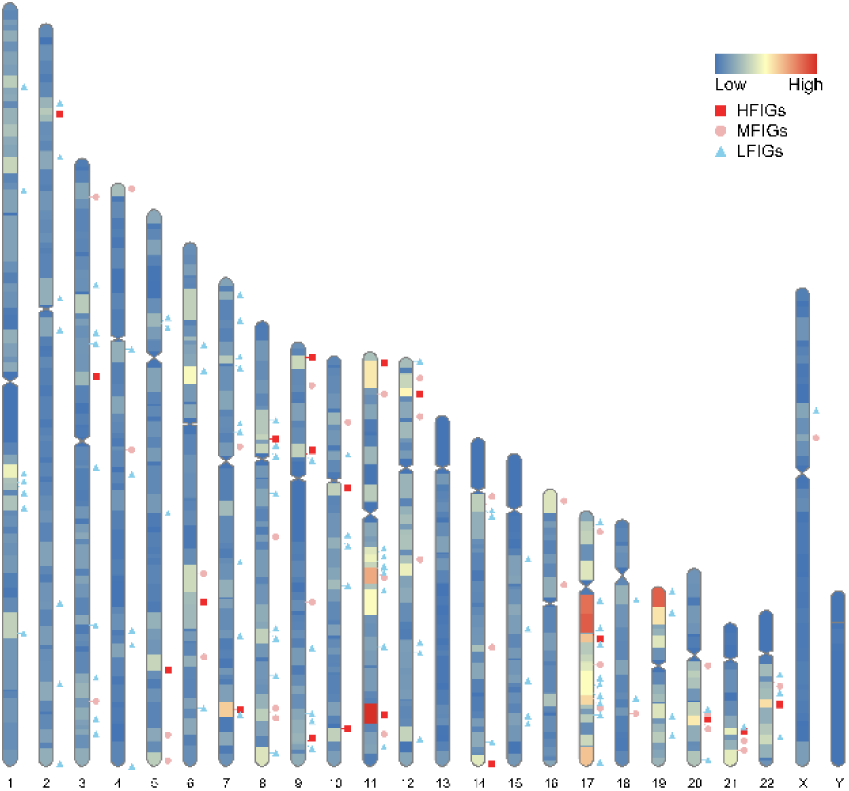
The distributions of interaction dense genes on chromosomes. The darker the color, the higher the number of banding interactions, and different shapes indicate different genes. HFIIGs (High frequency in genes), MFIIGs (Medium frequency in genes) and LFIIGs (Low frequency in genes) represent high interaction, medium interaction and low interaction genes respectively.

### Molecular mechanism. The repeat element impact

Approximately 45-60% of the human genome comprises repetitive sequences. Dynamic variations in the genomic positioning and copy number of these repeats may contribute to gene fusion events. We analyzed the repeat content near normal and fusion genes and observed no significant difference between the two groups (Fig.S1). We further analyzed and compared the content of various types of repeats, including SINEs, LINEs, Satellites, DNA transposons, simple_repeats, and LTRs, near fusion genes and normal genes. We found that, except for simple_repeats, there were significant differences in the distribution of the other repeat types near fusion genes and normal genes (Fig. S2). Consequently, we focused on a specific class of segmental duplications (SDs)-to further investigate their potential role in mediating gene fusion. SDs, defined as repetitive genomic regions exceeding 1 kb in length with >90% sequence similarity, exhibit high sequence homology and structural complexity, rendering them among the most mutation-prone genomic regions. Their inherent susceptibility to non-allelic homologous recombination (NAHR) frequently drives chromosomal structural variations, including deletions, inversions, and translocations. To explore the relationship between SD content and fusion gene formation, we quantified and compared the SD content proximal to fusion genes and normal genes. Across three distinct extension ranges, the SD content near normal genes has the smallest median, nearly approaching zero, when compared with the other three groups of fusion genes. This indicates that the SD content near fusion genes is higher than that near normal genes (Fig.7). Moreover, there is an extremely significant difference in SD content between normal genes and fusion genes (Fig.7). Given the established role of chromosomal structural variations in fusion gene generation and the propensity of SDs to induce such instability, the elevated SD density proximal to fusion genes strongly suggests a mechanistic association. Specifically, the enrichment of SDs near fusion gene loci likely predisposes these regions to genomic breakage and subsequent recombination events, thereby promoting the formation of fusion genes through structural rearrangements.

**Figure 7.**
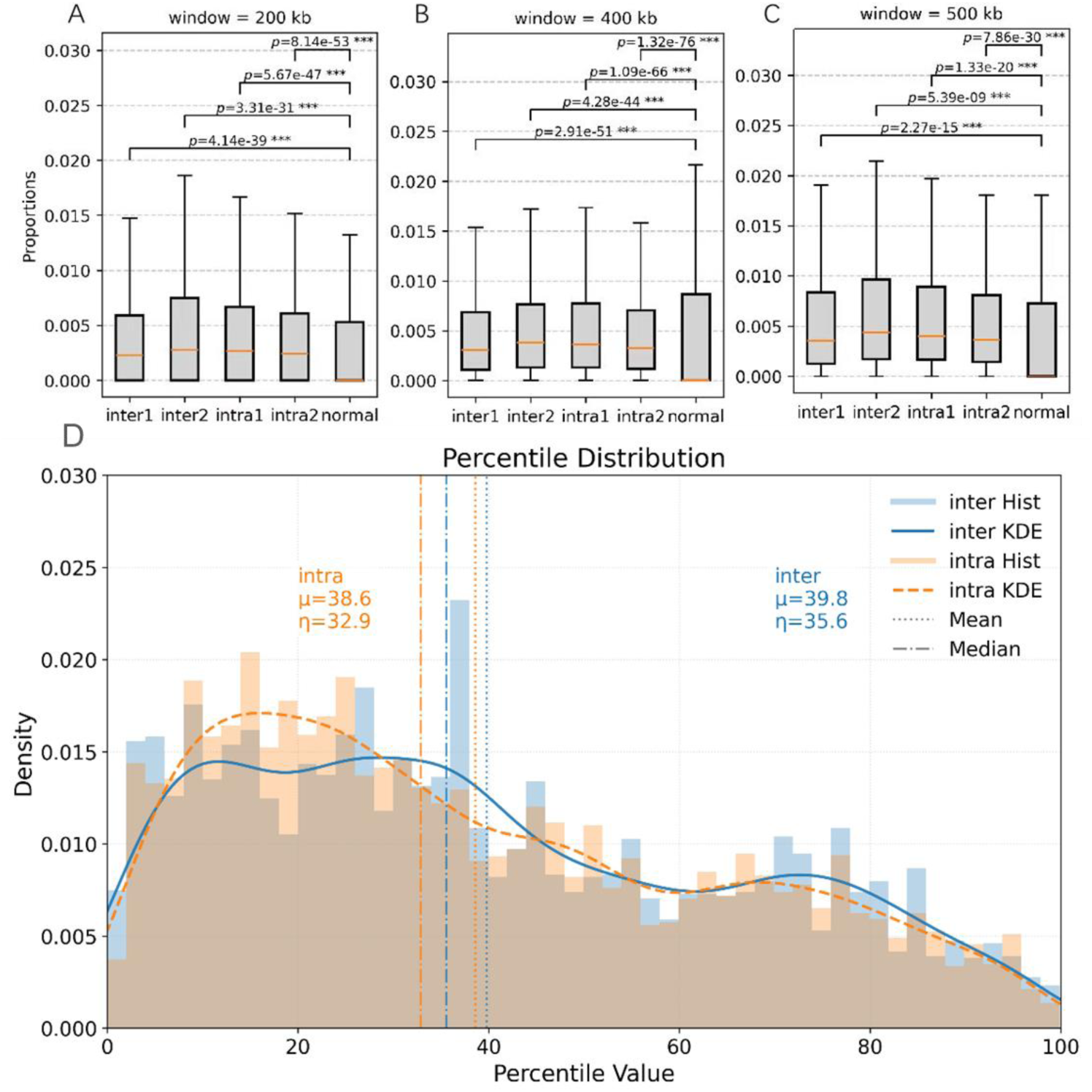
Distribution of SD content (A-C). The figure illustrates the distribution of SD content in fusion genes and normal genes across three extension ranges (500 kb, 400 kb, and 200 kb). In the figure, "inter" denotes intergenic interactions, "intra" represents intragenic interactions, and labels "1" and "2" correspond to the two genes participating in the fusion event. Red lines indicate median values, while the extension ranges (e.g, 500 kb) refer to the upstream and downstream regions flanking the original gene length. Asterisks (***) highlight statistically significant differences (p < 0.001) in SD content between fusion and normal genes. Notably, the median SD content near normal genes consistently approaches zero across all extension ranges, whereas fusion genes exhibit markedly elevated SD levels, underscoring the pronounced disparity between these groups. This observation aligns with the mechanistic role of SDs in promoting chromosomal instability and non-allelic homologous recombination, which are critical drivers of fusion gene formation. Percentile Distribution of GC Content Differences (D). The figure illustrates the percentile distributions of GC content differences for intergenic interactions (blue) and intragenic interactions (red) in fusion gene pairs. The distributions, represented by histograms combined with kernel density estimates, demonstrate distinct left-shifted peaks for both interaction types. For intergenic interactions, the mean (μ) and median (η) values were 39.4 and 37.2, respectively, while for intragenic interactions, they were 36.7 and 35.8. Both the median and mean values for all interactions fell below 40, indicating that the majority of percentiles clustered toward lower values. This pattern further supports the observation that the GC content differences between fusion gene pairs are significantly smaller than those between fusion genes and normal genes, emphasizing the role of GC content similarity in driving gene fusion events.

### GC content

The base composition of genes, particularly GC content, is also a key focus in studying the mechanisms of gene fusion. Meneu found that when the GC content of exogenous DNA was similar to that of the host genome, the host chromatin was more likely to accept the exogenous DNA (26). We propose that the GC content between fused gene pairs also exhibits similarity. So we investigated the similarity in GC content between fusion gene pairs. To analyze this, we calculated the GC content difference between a pair of fusion genes (|Di|) and the GC content difference between one of the fusion genes and all normal genes (|Eij|). We then determined the percentile rank of |Di| within the corresponding |Eij| distribution (eg., if 95% of the values were greater than |Di|, the percentile rank was 5%). We find that the percentile ranks for both types of interactions are skewed toward lower values, with medians and means below 40. This indicates that most percentile ranks are relatively small, suggesting that the GC content differences between fusion gene pairs are smaller than those between fusion genes and normal genes (Fig.7D). These results suggest that GC content similarity between gene pairs may facilitate fusion events.

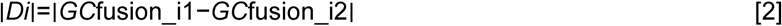

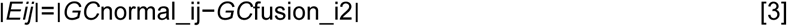

## Discussion

In this study, we selected blood disorders and solid tumors and compared them with Hi-C data (GM12878) in normal physiological states. Fig. 8 systematically showed the correlation analysis of Hi-C data and fusion gene events on chromosome 17. It could be obviously seen that there was a significant spatial overlap between the Hi-C interaction matrix and the gene fusion event matrix (70), accounting for 94.59% of the total frequent regions of the fusion gene. The results showed that the regions with higher contact intensity in the three-dimensional space of chromosomes tended to have more fusion gene events. After that, the correlation between Hi-C signal, fusion event frequency and genome distance of banding was analyzed. The normal physiological cells and fusion genes had consistent tendency, that is, the interaction frequency decreased with the increase of distance, which further demonstrated that fusion gene events were liable to occur in spatially close segments. Finally, the Hi-C interaction intensity distribution between the fusion region and the non-fusion region was investigated. Compared with the non-fusion region, the Hi-C in the fusion region had a wider interaction intensity range and higher interaction intensity, and contained significantly more high Hi-C signal events. All above results fully proved the high correlation between fusion gene events and the spatial hot spots in the three-dimensional structure of chromosomes, and also confirmed the feasibility of determining the three-dimensional structure of chromatin based on fusion genes (Fig. S3).

**Figure 8.**
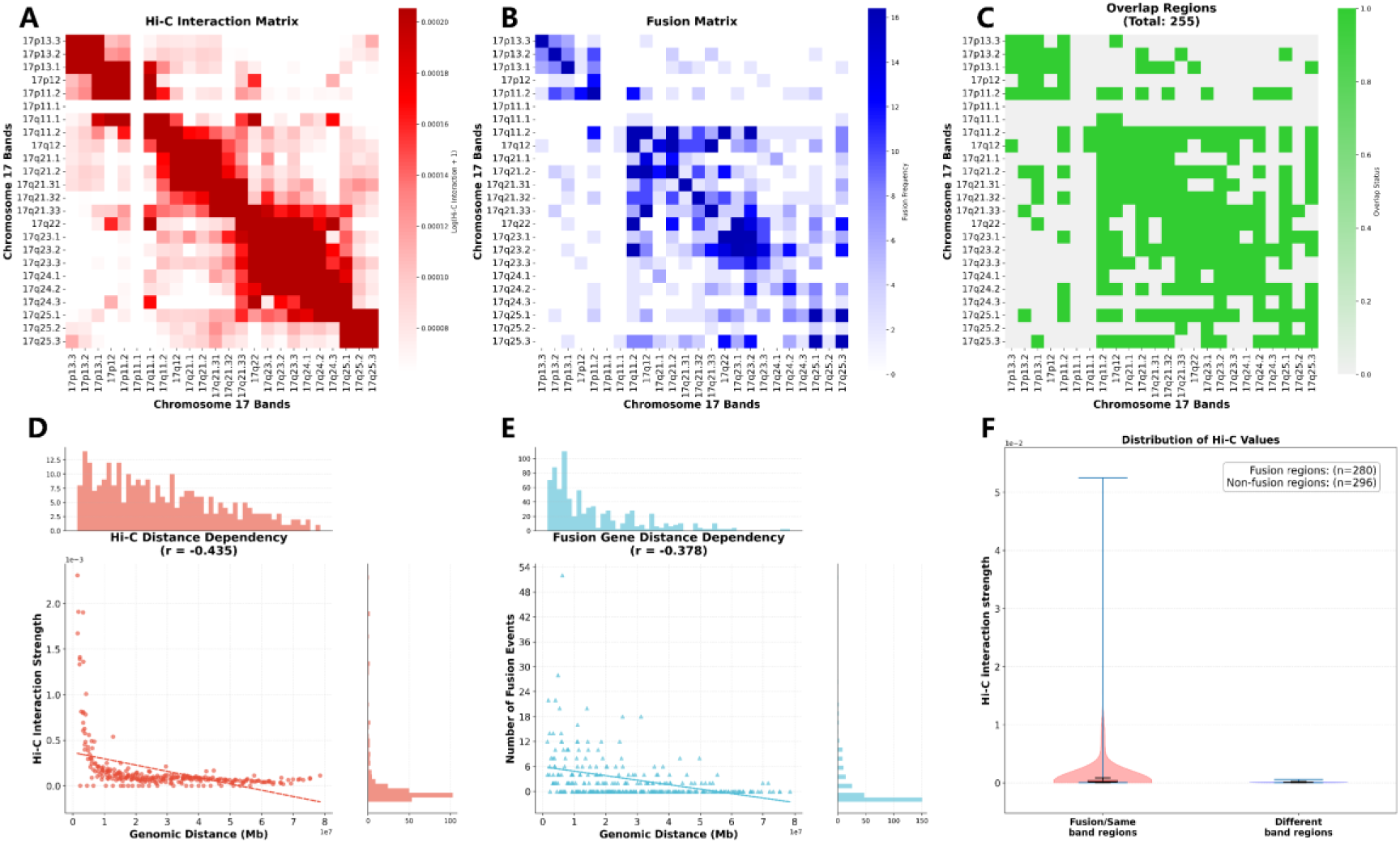
Comparison of Hi-C data and fusion gene data on chromosome 17. (a) and (b) are respectively the heat maps of the Hi-C normal human cell GM12878 cell line aggregated by banding and heat maps based on fused gene disease data. (c) represents the overlapping part of the heat map of Hi-C and the fusion gene. (d-e) are respectively the distance decay curves of Hi-C and the fusion gene. (f) describes the comparison of fusion genes and Hi-C data.

Chromosome 11 is a hot spot for structural variation, *KMT2A* on it is one of the most common mutated genes in hematopoietic malignancies and solid tumors. The unique genomic features of *KMT2A*, including long gene span, repeat enrichment, broken cluster region, and location at the classic fragile site *FRA11G*, make it prone to chromosome instability under DNA replication stress [27]. After passing through the DNA double-strand break repair pathway, *KMT2A* will misconnection to partner genes to form fusion proteins, and lose normal methylation function, which directly leads to abnormal expression of *KMT2A* target genes (HOX) [28]. On the other hand, *KMT2A* fusion proteins alter chromatin accessibility by binding to enhancer regions and recruiting various abnormal epigenetic regulators, transcription extension complexes, and chromatin remodeling factors. At the same time, *KMT2A* mediates the formation of abnormal enhancer-Hoxa rings, locking in the continuous activation state of oncogenes [29]. In this state of three-dimensional structural disorder, *HOX* will continue to overexpress or abnormally silence, driving the malignant transformation of hematopoietic malignant tumors and solid tumors. She et al. reported another possible mechanism, *KMT2A* mediates the K143 methylation of Borealin, and fusion of *KMT2A* can alter the localization of chromosome passenger complex in the inner centromere, as well as liquid-liquid phase separation, increasing the risk of cancer [30].The molecular mechanism of ETV6 to form fusion genes is similar to *KMT2A*. *ETV6* is rich in long introns and repetitive sequence, located in the genome vulnerable site FRA12C, easy to break under replication pressure [31]. After the fusion event, the C-terminal inhibitory domain (*HLH*/*POZ* domain) of *ETV6* failed, and the abnormal activation domain would be obtained after fusion with *RUNX1*, *ABL1*, *NTRK3*, permitting the fusion protein carcinogenicity [32, 33]. While N-terminal PNT domain would hijack the target gene promoter leading to its continued activation [34], on the other hand, recruit transcription complex, kinase signaling complex and chromatin remodeling complex to drive carcinogenic signaling pathways.Abnormal connections in the V(D)J recombination region of *IGH* are the key inducement of fusion events [35]. The regulatory region of *IGH* performs hypomethylated in B cells and causes open chromatin. Transcription factors such as PAX5 and EBF1 bind with mediators to activate high transcriptional activity. Super-enhancer of *IGH* Eμ hijacks oncogenes and recruits both *SWI*/*SNF* and RNA polymerase II, leading to persistent oncogene expression [36–38]. 3’RR binds to oncogene promoters via chromatin loops, recruits *BRD4* and *P-TEFb*, and activates RNA polymerase II extension activity to drive oncogene expression [39–41].

Our fusion gene technology has several key advantages in studying the three-dimensional organization of chromatin. Our method is essentially an economical and efficient technical alternative that can achieve high-throughput analysis of a large number of samples. The conservation of the same chromosomal spatial conformation among different cancer types was verified by using the derived data from genomic sequencing. Meanwhile, this method avoids complex experimental procedures and is compatible with routine clinical samples. In addition, this study achieved re-annotation of traditional genomic data from a three-dimensional mechanism, directly associating the distribution of chromosomal translocations with driver gene fusions, providing a mechanism basis for targeted therapy. Our chromosome localization method based on fusion genes also has its own limitations. Firstly, our method has a relatively high resolution (in units of bandings), lacks normal sample controls, and the conclusion is a probabilistic model, which cannot precisely locate the chromosomal coordinates of individual tumor cells. If multi-omics techniques (such as single-cell Hi-C) can be combined, more accurate chromosomal spatial conformations can be obtained, and the association between chromosomal dynamic recombination and treatment response can be revealed (42).

This study, through the analysis of a large amount of trans-chromosomal interaction data, revealed the universality of chromosomal spatial conformations in cancer, providing new ideas for the development of diagnostic tools and targeted strategies. The results indicated that different cancer types shared the same chromosomal spatial arrangement pattern, and pan-cancer diagnostic markers based on chromosomal spatial localization could be developed. Abnormal nuclear structure is a pathological marker of malignant tumors [43, 44]. We have found that high-frequency translocation chromosomes are usually located in the nuclear center and are prone to gene fusion. Therefore, by analyzing the distribution pattern of chromosomal translocations within the cell nucleus of tumor cells, it is possible to assist in identifying nuclear structural abnormalities in malignant tumors. In addition, genes carried by hub chromosomes can be prioritized as candidate targets for targeted therapy or gene editing (e.g., KMT2A, NUP98, RARA). The development of anti-translocation drugs is also a feasible approach. Inhibiting the error repair of "translocation vulnerable areas" can selectively kill tumor cells [45, 46].

## Materials and Methods

### 1. Fusion gene retrieval and analysis

Using the web site https://atlasgeneticsoncology.org, https://www.kobic.re.kr/chimerdb/, https://cancer.sanger.ac.uk/cosmic, https://mitelmandatabase.isb-cgc.org/ for retrieval of fusion gene, find 23 chromosomes in blood diseases and fusion gene in solid tumors [30]. Combined with NCBI (https://www.ncbi.nlm.nih.gov/), to determine chromosome banding, gene starting position and end position of the information.

### 2. Data analysis and mapping

For the data of gene fusion of different chromosomes in blood diseases and solid tumors, SPSS 22 statistical software (SPSS Inc., IL, USA) and Excel 2010 were used for processing and analysis, and the data were sorted into trans-chromosome interaction and cis-chromosome interaction. Software such as python, R, etc. was used to draw the chord diagram of the trans-chromosome interaction, and the code used to generate the results and diagrams from the above analysis is available from the author upon request.

### 3. Integrating gene data organization and plotting the three-dimensional structure of chromosomes

Converting the fusion gene database file into a fusion gene summary file and perform de-duplication organization. Preparing the chromosome banding file including band, start, and end. Using the script find_gene.py to compare the gene name with the genomic.gff parsing file in the ncbi database and combine it with chromosome banding localization to obtain Fusion_pair. gene1, gene1 band, gene1 start, gene1 end, gene1 length, gene2, gene2 band, gene2 start, gene2 end, gene2 length. The number of occurrences of the same gene1 band and gene2 band data statistics is recorded as frequency, and they are arranged in the order of the reference genome banping to obtain the contact_frequency.csv file with three columns as band1, band2, and frequency respectively.

The three-dimensional structure of the chromosome was constructed using the pastis_NB algorithm with banded resolution. Since it is a sparse matrix of 862*862 and the data was overly discrete with variance greater than the mean, negative binomial statistics were adopted and the script pm2_reconstructions.py was executed. This script sets the expected contact frequency *μ_ij_* between the visible bands as the exponential negative correlation function of their Euclidean distance in three-dimensional space

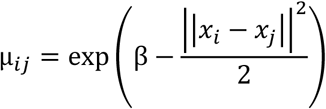

among them, *x_i_* and *x_j_* represent the three-dimensional coordinates of the bands i and j respectively. Subsequently, the three-dimensional coordinates of all banders are solved by maximizing the log-likelihood function based on the negative binomial distribution.

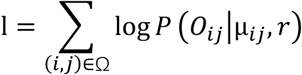

Visualizing the generated file of PM2.structure.connected.pdb, this file contains Chr, band, x, y, z five columns, the file is the space coordinates of banding. Using the script plot_total_chr.py to call the matplotlib visualization structure of python.

### 4. SD content and GC content

This study adopted the hg38 reference genome and annotation file (26). Convert the fusion gene list file into a bed file, divide it into four groups, and merge them simultaneously to obtain the all-fusion.bed file. Extract all the genetic information in the annotation file of hg38 to obtain the bed files of all normal genes. Extend the fused genes within a certain range before and after, with the extension range being a variable, to obtain the bed file after the extension range of the fused genes. Use the same operation for the four groups of fused genes. Apply the above operation to the normal gene to obtain the extended normal gene bed file, and remove the parts that repeat with the fused gene in the extended normal gene file. Use the command "bedtools coverage" for the fusion gene bed file and the SD_bed file to obtain the overlapping range and size between the two files, as well as the proportion of SDs. The same operation was applied to the four groups of fused genes and normal genes, and the SDs contents within the extension range of the four groups of fused genes and normal genes were obtained respectively. Using the same method, the differences in the overall repeats content and the differences in the contents of multiple types of repeats between the fused gene and the normal gene were compared.

## Acknowledgments

This work was supported by the National Key Research and Development Program of China (Grant No.2022YFC3401300), the National Natural Science Foundation of China (Grant No. 32471540, 32371540, 21878128), Wuxi City’s "Double Hundred" Young and Middle aged Medical and Health Top Talents: BJ2023004, Jiangsu Province Preventive Medicine General Project: Ym2023008, and Reserve Discipline Leader of Wuxi People’s Hospital: 2024-YZ-HBDTR-WQ-2024.

## Author Contributions

H.-L.Y. and Q.W. designed and funded; Y.-Q.C. wrote draft and edited; X.-Z.R. and Y.-L.D. conducted the calculations; C.-T.Y. designed frame and edited. Z-Y.H. helped with chord diagram; J.-A.Y. helped with editing; S.-S.F. analyzed data; L.-F.L. supported the Hi-C analysis.

