## Supplemental S1-S3 for "The fusion gene occurrence spectrum predicts the spatial structure of chromosomes"

**This PDF file includes:**

Supporting text

Figures. S1 Distribution of repeats content.

Figures. S2 Content of various types of repeats.

Figures. S3 Comparison of Hi-C data and fusion gene data on chromosome 19, 20, 22 from top to bottom.

**Supporting Information Figures**

Fig. S1. The figure illustrates the distribution of repeat content in fusion genes and normal genes across three extension ranges (500 kb, 400 kb, and 200 kb). In the figure, "inter" denotes intergenic interactions, "intra" represents intragenic interactions, and labels "1" and "2" correspond to the two genes participating in the fusion event. Red lines indicate median values, from the figure, it can be seen that under three types of extension ranges, the medians of repeats content for fusion genes and normal genes are roughly the same, around 0.5. However, there are differences in the upper and lower limits of the boxes and the extreme values.

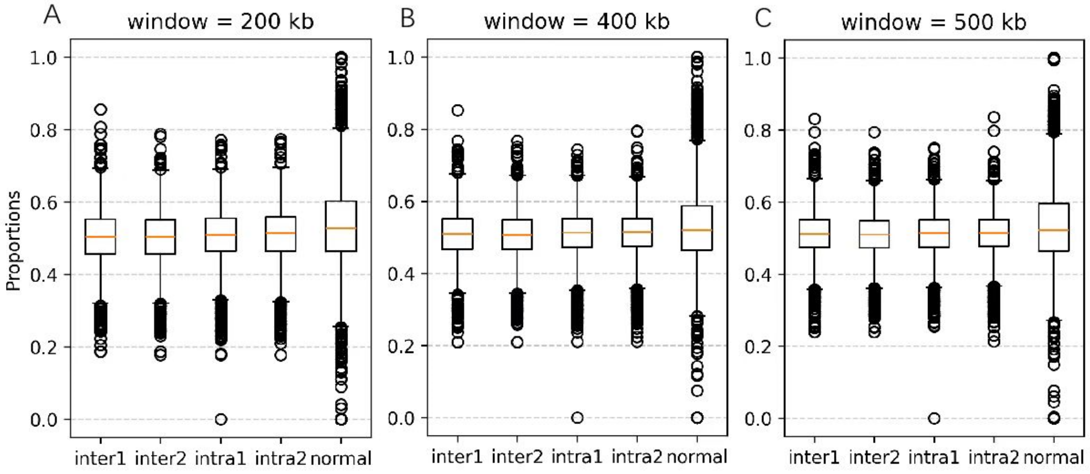

Fig. S2. These figures illustrate the distribution of various type of repeat content in fusion genes and normal genes across one extension ranges (500 kb). In the figure, "inter" denotes intergenic interactions, "intra" represents intragenic interactions, and labels "1" and "2" correspond to the two genes participating in the fusion event. Red lines indicate median values, asterisks (***) highlight statistically significant differences (p < 0.001) in SD content between fusion and normal genes. From the figure, it can be seen that for normal genes and fusion genes, the distribution of repeat content varies with different types of repeats.

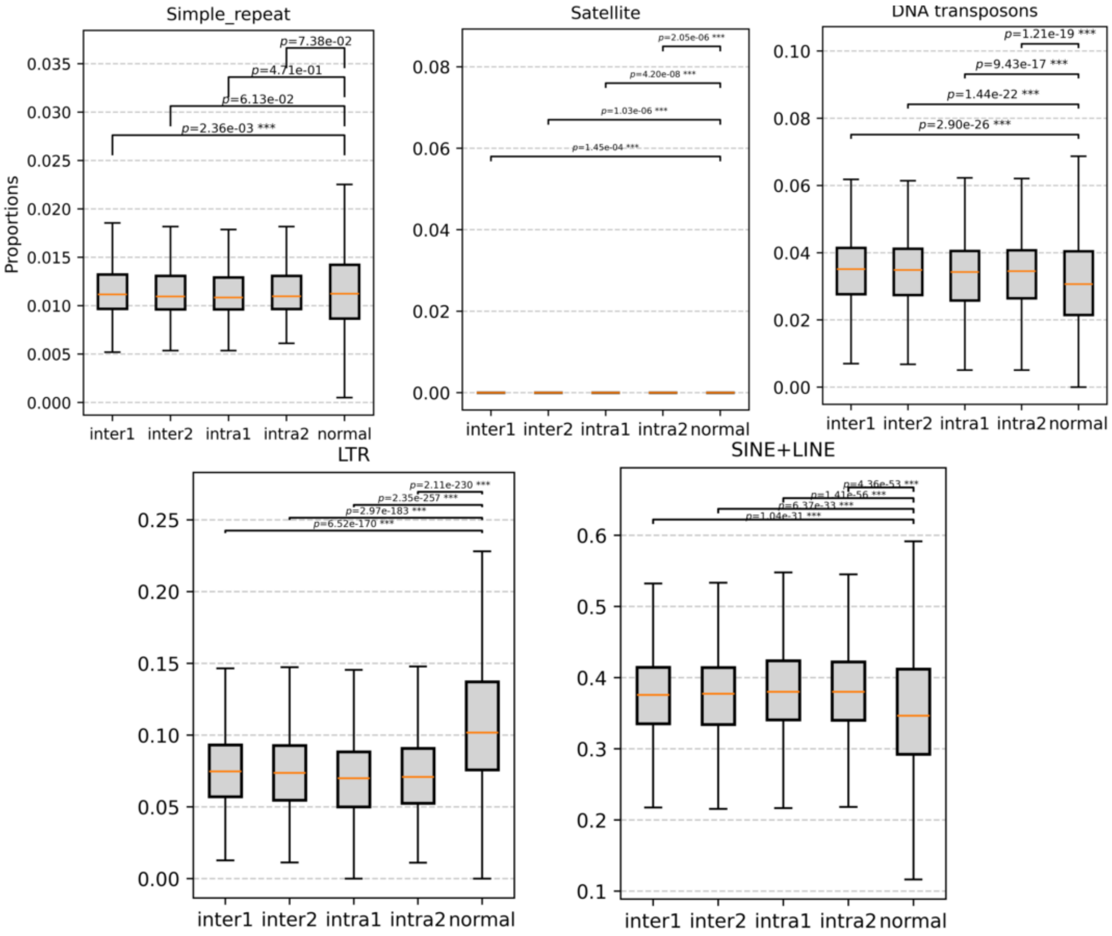

Fig. S3. (a) and (b) are respectively the heat maps of the Hi-C normal human cell GM12878 cell line aggregated by banding and heat maps based on fused gene disease data. (c) represents the overlapping part of the heat map of Hi-C and the fusion gene. (d-e) are respectively the distance decay curves of Hi-C and the fusion gene. (f) describes the comparison of fusion genes and Hi-C data.

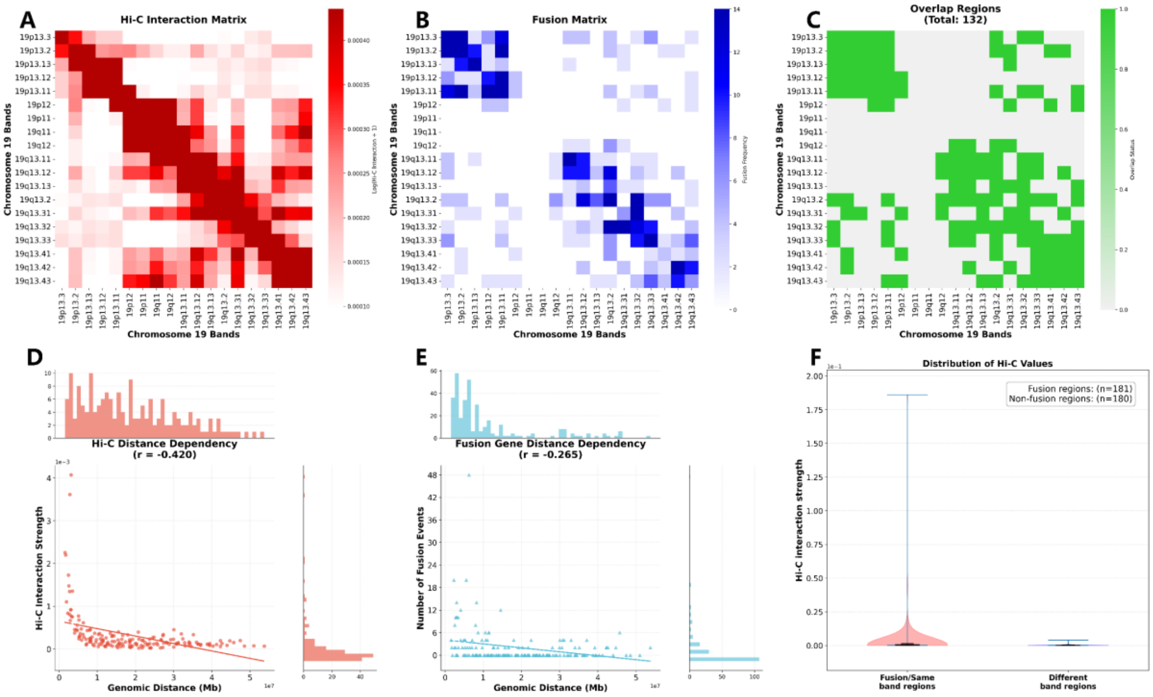

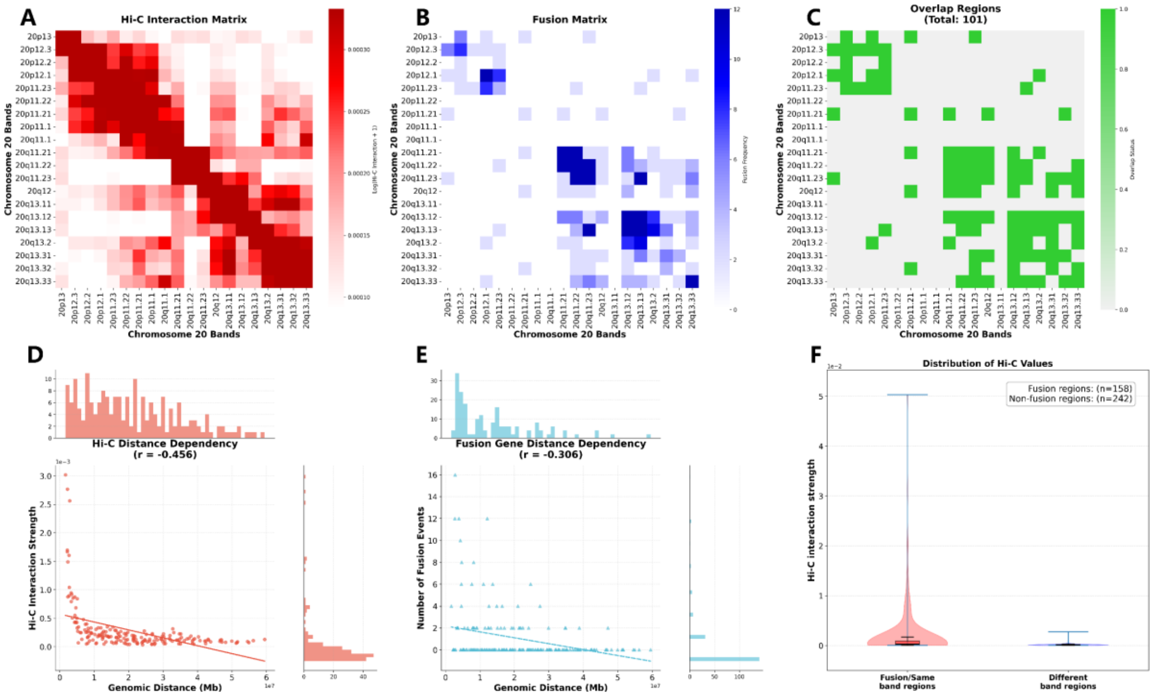
A
Hi-C Interaction Matrix
Chromosome 20 Bands
B
Fusion Matrix
C
Overlap Regions
(Total: 101)
D
Hi-C Distance Dependency
(r = -0.456)
Hi-C Interaction Strength
Genomic Distance (Mb)
E
Fusion Gene Distance Dependency
(r = -0.306)
Number of Fusion Events
F
Distribution of Hi-C Values
Fusion regions: (n=158)
Non-fusion regions: (n=242)
Hi-C interaction strength
Fusion/Same band regions
Different band regions

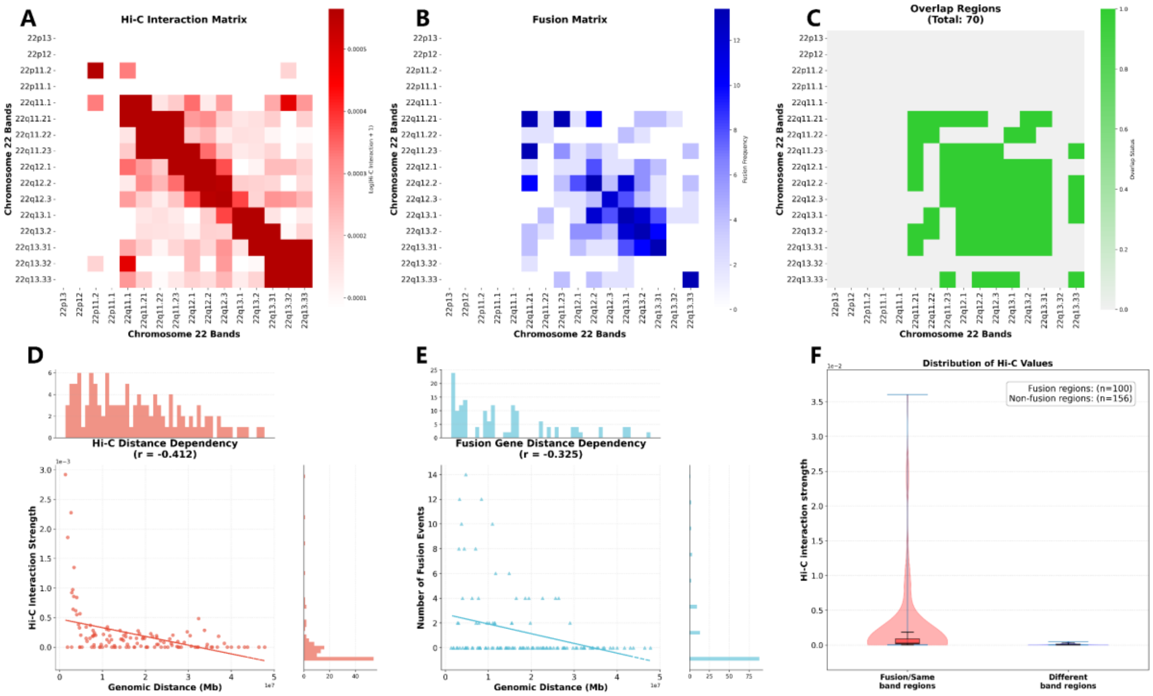
A
Hi-C Interaction Matrix
Chromosome 22 Bands
B
Fusion Matrix
C
Overlap Regions
(Total: 70)
D
Hi-C Distance Dependency
(r = -0.412)
Hi-C Interaction Strength
Genomic Distance (Mb)
E
Fusion Gene Distance Dependency
(r = -0.325)
Number of Fusion Events
F
Distribution of Hi-C Values
Fusion regions: (n=100)
Non-fusion regions: (n=156)
Hi-C interaction strength
Fusion/Same band regions
Different band regions
